# Applications of some artificial intelligence tools in the drug design of some compounds targeting the main viral protease of the Feline Infectious Peritonitis Virus (FIPV) in cats

**DOI:** 10.1101/2024.09.09.612092

**Authors:** Mohd Yasir Khan, Abid Ullah Shah, Nithyadevi Duraisamy, Nadine Moawad, Reda Nacif ElAlaoui, Mohammed Cherkaoui, Maged Gomaa Hemida

**Affiliations:** Department of Computer Science, College of Digital Engineering and Artificial Intelligence, Long Island University, Brooklyn, (MYK), (ND), (RNE), (MC); Department of Veterinary Biomedical Sciences, College of Veterinary Medicine, Long Island University, 720 Northern Boulevard, Brookville, NY, 11548, USA, (AS), (NM), (MGH)

**Keywords:** FIPV, M^pro^, In silico design, molecular docking, antiviral

## Abstract

Feline infectious peritonitis virus (FIPV) is one of cats’ most serious viral infections. The FIPV infection induces a complicated syndrome in the affected cats, including immunosuppression and severe inflammatory conditions. Unfortunately, these vaccines cannot prevent cats from getting infected with these viral infections. There is ongoing research on preparing antiviral therapies against FIPV in cats. However, these are still in clinical trials and have not been fully approved by the drug authorities in many countries, including the USA. Targeting the main viral proteases is one of the promising trends in the drug design of many viral diseases, including coronaviruses. The main goal of the current study was to repurpose and test the efficacy of some known antiviral drugs to treat FIPV infection in cats by targeting the FIPV-main protease enzyme. To achieve these goals, we used the in-silico prediction and molecular docking tools to screen and identify some drugs targeting FIPV-MPro. We used the docking and binding energies as the main parameters for selecting target compounds (FIPV-MPro). Our results show that out of the 15 antiviral and immunomodulatory compounds, the top-ranked inhibitors for the FIPV-Mpro are (Michael acceptor inhibitors (N3), Sofosbuvir, and methotrexate).In conclusion, our results confirmed the potential applications of the predicted FIPV-Mpro inhibitors either independently or in combination with other immune-modulatory compounds. Further in vitro and in vivo studies are encouraged to test the efficacy of these identified compounds as potent inhibitors for the MPro of the FIPV in cats. This study will pave the way for the development of novel drugs that treat FIPV infection in cats.

## 1 Introduction

FIPV belongs to the order Mononegavirales, family coronaviruses and genus alphacoronavirus, species alphacoronavirus-1 and subspecies feline coronavirus (FCoV). The viral genome is a single molecule of a positive sense RNA. The FIPV genome has a typical coronavirus genome organization and encodes 11 proteins, including four structural and seven non-structural proteins. The major non-structural proteins encoded by Gene-1 (ORF1a and ORF1b with ribosomal frameshifting in between) reside at the 5’ two-thirds of the genome. Other non-structural proteins of the FCoV are encoded by the 3 abc and 7a/b genes. While the structural proteins (Spike (s), Envelope (E) Membrane (M), and the nucleocapsid (N)). The ORF1a/b is further processed by some viral encoded proteases into 16 non-structural proteins (NSP-1-NSP16) [1]. The FIPV-3c protein is important for virus replication and contributes substantially to viral virulence and tissue tropism [2]. Although the function of the FIPV-7ab proteins is not fully understood, they might play important roles in viral immune evasion, particularly as an antagonist for the IFN-type I [3]. Based on the FIPV-S sequences, the virus is classified into two serotypes. The serotype-I is designated as FIPV, which is the most virulent serotype and causes a lethal infection in cats [4]. The serotype II is considered the feline enteric coronavirus FEC, which usually induces asymptomatic or mild infections in cats [4]. Although FIPV infection in cats is not contagious (it does not spread easily among cat populations) until now, the prognosis of the cats infected with the virus is always fatal in most cases [5]. The absence of vaccines that could protect cats against FIPV infection makes antiviral therapy the only remedy to treat cats from FIPV infection. Several antiviral compounds, particularly nucleoside analogs and interferons, have been tried to inhibit or interrupt the FIPV replication cycle at various stages [6–9]. This approach includes the application of a single or a combination of different compounds together to ensure the robust inhibition of the viral replication [10]. Coronaviruses main protease (Mpro) is one of the most important targets for the design of antiviral therapy for several coronaviruses, including the Middle East Respiratory syndrome coronavirus (MERS-CoV) and the Sever acute-2 (SARS-CoV-2), the porcine epidemic diarrhea virus (PEDV) as well as the FIPV [11]. The main reason for targeting CoVs-Mpro is their essential role in polyprotein processing during viral replications. In the current study we used the in-silico drug design tools to predict the efficacy of 15 selected antiviral, anti-inflammatory and immune-modulatory compounds on the inhibition of the FIPV-Mpro enzyme (the Michael acceptor inhibitor (N3N3) compound, Oxipurinol, Favipiravir, Pentoxifylline, Baricitinib, Methotrexate, Gemcitabine, Galidesivir, Ribavirin, Azauridine, GS441524, Mizoribine, Sofosbuvir, Molnupiravir, and Tenofovir). We used the crystal structure of the FIPV-Mpro available in the public domain and analyzed the binding sites of each compound to these in the FIPV-Mpro. This approach will provide more insights into the inhibitory actions of the selected compounds on the FIPV-Mro. These potential FIPV drugs could be used independently or in combination to inhibit FIPV replication, to calm down the severe inflammatory conditions associated with the viral infections, and to enhance the immune response against the viral infection. It will also pave the way for more research to do more functional characterization of the potential inhibitory effects of these compounds on the FIPV replication in the in vitro and in vivo models. This approach will lead to the production of effective antiviral therapy against FIPV infection in cats.

## 2 Materials and Methods

### 2.1. Retrieval of the FIPV major proteins and ligands

The FIPV main protease (FIPV-Mpro) protein in complex with ligand N3 and their 3D structures were downloaded from the RCSB-Protein Data Bank (PDB) (https://www.rcsb.org/), in the PDB file [12]. The PDB ID of FIPV-Mpro is 5EU8. Further, all other ligands (compounds) used in this study were retrieved from the PubChem database https://pubchem.ncbi.nlm.nih.gov. The PubChem compound ID (CID) (https://pubchem.ncbi.nlm.nih.gov/) for each ligand is given in (Table 1).

**Table 1:** Summary of the molecular docking data of the selected 15 compounds with the FIPV-Mpro, including the binding energy and CDOCKER scores.

| N | Ligand PubChem CID | Ligand name | Binding Energy ($\Delta G$) (kcal/mol) | CDOCKER Energy Score |
| --- | --- | --- | --- | --- |
| 1 | 6323191 | N3<br>(bound reference ligand) | -246 | -78.82 |
| 2 | 45375808 | Sofosbuvir | -232.76 | -46 |
| 3 | 126941 | Methotrexate | -229.42 | -44.55 |
| 4 | 145996610 | Molnupiravir | -202.54 | -12.96 |
| 5 | 10445549 | Galidesivir | -201.36 | 1.80 |
| 6 | 464205 | Tenofovir | -185.24 | -30.53 |
| 7 | 44205240 | Baricitinib | -162.05 | 25.79 |
| 8 | 37542 | Ribavirin | -161.18 | 15.23 |
| 9 | 60750 | Gemcitabine | -157.37 | -9.6 |
| 10 | 104762 | Mizoribine | -152.50 | -7.10 |
| 11 | 44468216 | GS441524 | -151.54 | 4.08 |
| 12 | 5901 | 6-Azaauridine | -148.33 | 3.22 |
| 13 | 4740 | Pentoxifylline | -130.82 | -21.57 |
| 14 | 492405 | Favipiravir | -115.58 | -8.8 |
| 15 | 135398752 | Oxipurinol | -101.3 | -4.37 |

### 2.2. Preparation of the target proteins and ligands

For the docking process, the protein molecules and compounds as ligands need to be prepared using the protein and ligand preparation step by Biovia Discovery Studio v22.1.021297. This process typically involves tasks like removing water molecules and other heteroatoms, adding hydrogens, and energy-minimization to ensure accurate molecular docking as previously described [13].

### 2.3. Parameters for the prediction of the protein’s active sites

The active site is the target for an enzyme’s inhibition. In active site prediction, the bound reference ligand or test ligand was used as a probe for the active site prediction [14]. The active site was identified as amino acids within the 10 Å range of the bound reference ligand. The reference ligand binding site sphere attributes (X: -47.58 Y:-15.16 Z: -9.49) were used to dock ligands. Mapping the location of the binding site before the docking process significantly increases the docking efficiency. Also, one can obtain information about the binding sites by comparing the target protein with a family of proteins of interest sharing a similar function or with proteins co-crystallized with other ligands.

### 2.4. Molecular docking of the selected compounds with the FIPV-Mpro

The molecular docking approach can be used to model the interaction between the selected molecules and the target proteins. The interactions between these molecules and the proteins of interest at the atomic level of molecular docking revealed this interaction’s potential function and action and their involvement in various biological processes [15]. We typically use ten docking poses for ligands in the CDOCKER tools of the Biovia Discovery Studio (https://www.3ds.com/products/biovia/discovery-studio) for our molecular docking protocol. For each dock ligand pose, the higher the CDOCKER Energy scores’ negative values and the calculated binding energy (ΔG) indicate more favorable binding conditions [14, 16, 17]. These computational tools provide options for the visualization of the ligand-target interaction (molecular docking) and the identification of the compounds that bind more efficiently with the target proteins (2). This analysis typically involves the docking scores, the ligand-protein interactions, and the visualization of the docked complexes. To identify the binding sites within the target proteins, we calculated the binding energy (ΔG) as an internal validation and benchmarking step using the Biovia Discovery Studio v22.1.021297 as described earlier [14].

### 2.5 Analysis of the FIPV-Mpro-ligands complex

Protein-ligand interactions were analyzed, and illustrations were generated using the Biovia Discovery Studio visualizer. We used some criteria for the identification of the hydrogen bonds including two major specific parameters: (1) the distance between the donor (D) and acceptor (A) atoms must be 3.35 Å or less (with 3.35 Å being the maximum allowable distance), and (2) the distance between the hydrogen (H) and acceptor (A) atoms must be 2.70 Å or less (with 2.70 Å being the maximum allowable distance). In addition, the non-bonded contacts refer to the interactions between the hydrophobic atoms, such as the carbon (C) or the sulfur (S), and any other atoms that are neither covalently bonded nor involved in hydrogen bonding. The non-bonded interactions occur within a distance range of (2.9 to 3.9 Å) [18].

## 3. Results

### 3.1. FIPV-Mpro interaction with the N3 compound

The FIPV-MPro interaction with the N3 compound in the X-ray crystal structure retrieved from the RCSB-PDB database (PDB ID:5eu8; 1.80 Å resolution) was chosen as the standard for different ligands docking. The PubChem CID for all ligands is shown in (Table 1). The best docking pose of the N3/ FIPV-Mpro interaction indicated a strong binding affinity with the test compounds’ highest binding energy (ΔG= -246.66 Kcal/mol) (Table 1). However, the highest CDOCKER energy score (negative value) referred to the protein’s most favorable binding of ligands. The CDOCKER energy score for the same pose is (-**78.82)**. The interaction between FIPV-Mpro and N3 involving some key amino acid residues and bonds, including the Cys-144 (bond: Pi-Sulphur), the His-163, Glu-165, the Ser-189 and the Gln-191 (bond: Conventional hydrogen), the Leu-164 (bond: Alkyl and Pi-alkyl), the His-41, the Pro-188 and the Leu-166 (bond: carbon-hydrogen) shown in the 2D image (Figure 1) and (Table 1S).

**Figure 1:**
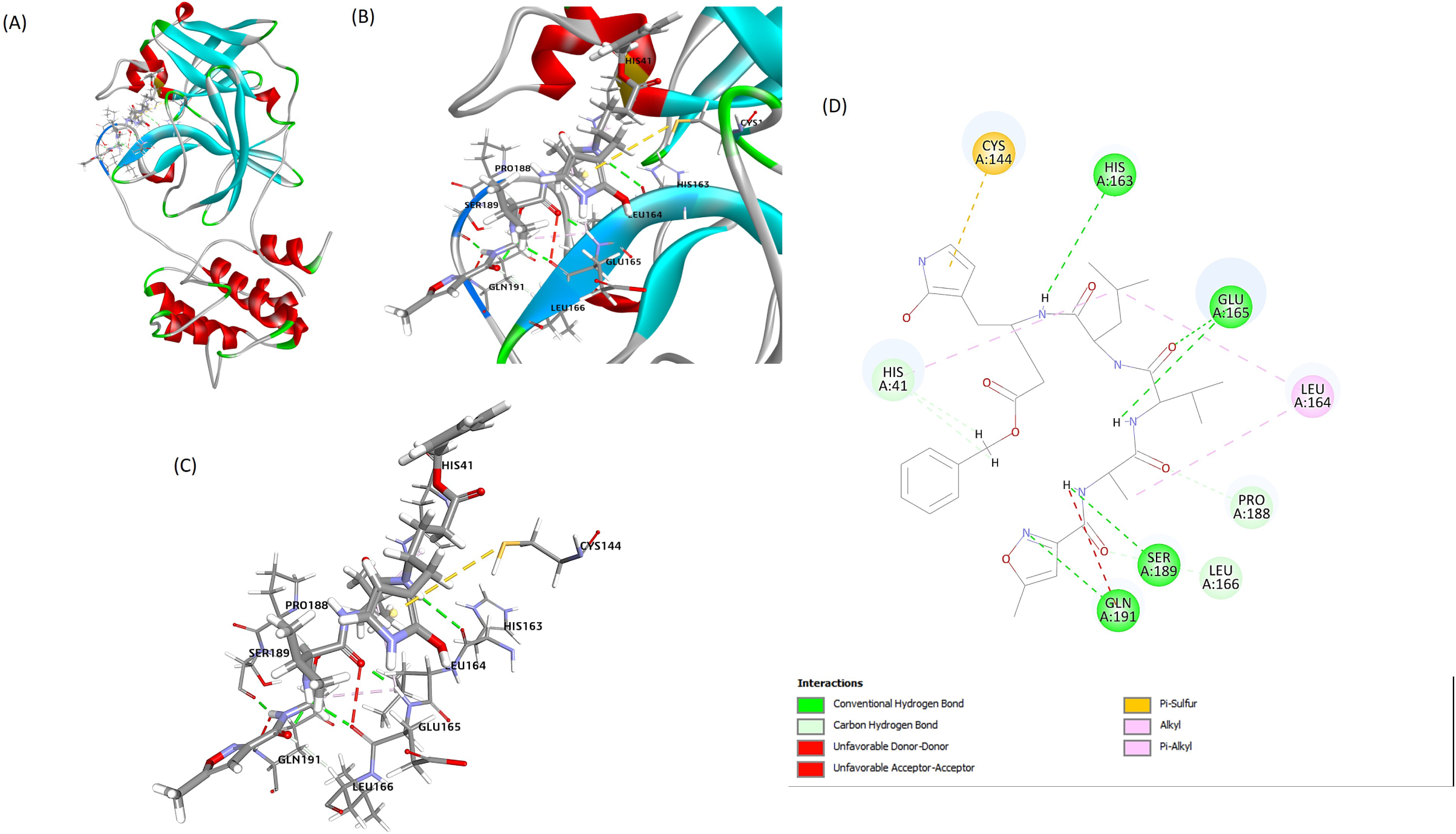
The best docking pose of the interaction of the FIPV-Mpro/ N3 interaction. (A) Mpro-N3 complex after docking (B) and (C) showing N3 interaction with residues of Mpro. (D) Interaction of N3 with residues including (CYS144, LEU164, MET190, GLU165, HIS41, HIS162, HIS163, SER189 and PRO188).

### 3.2. FIPV-Mpro interaction with the Oxipurinol

The best docking pose for the interaction between the FIPV-Mro and the **Oxipurinol** compound indicated the best binding affinity with the highest binding energy (**ΔG**= **-101.02** Kcal/mol) (Table 1). The CDOCKER energy score for the same pose is (**-4.37)**. The interaction between the FIPV-Mpro and the Oxipurinol involves some key amino acid residues and bonds, including Cys-144 (bond: Pi-Sulphur and hydrogen), Thr-47 (Bond: hydrogen), His-41 (bond: Pi-Pi T shaped), Pro-188 (bond: Pi-alkyl) shown in the 2D image (Figure 2) and (Table 1S).

**Figure 2:**
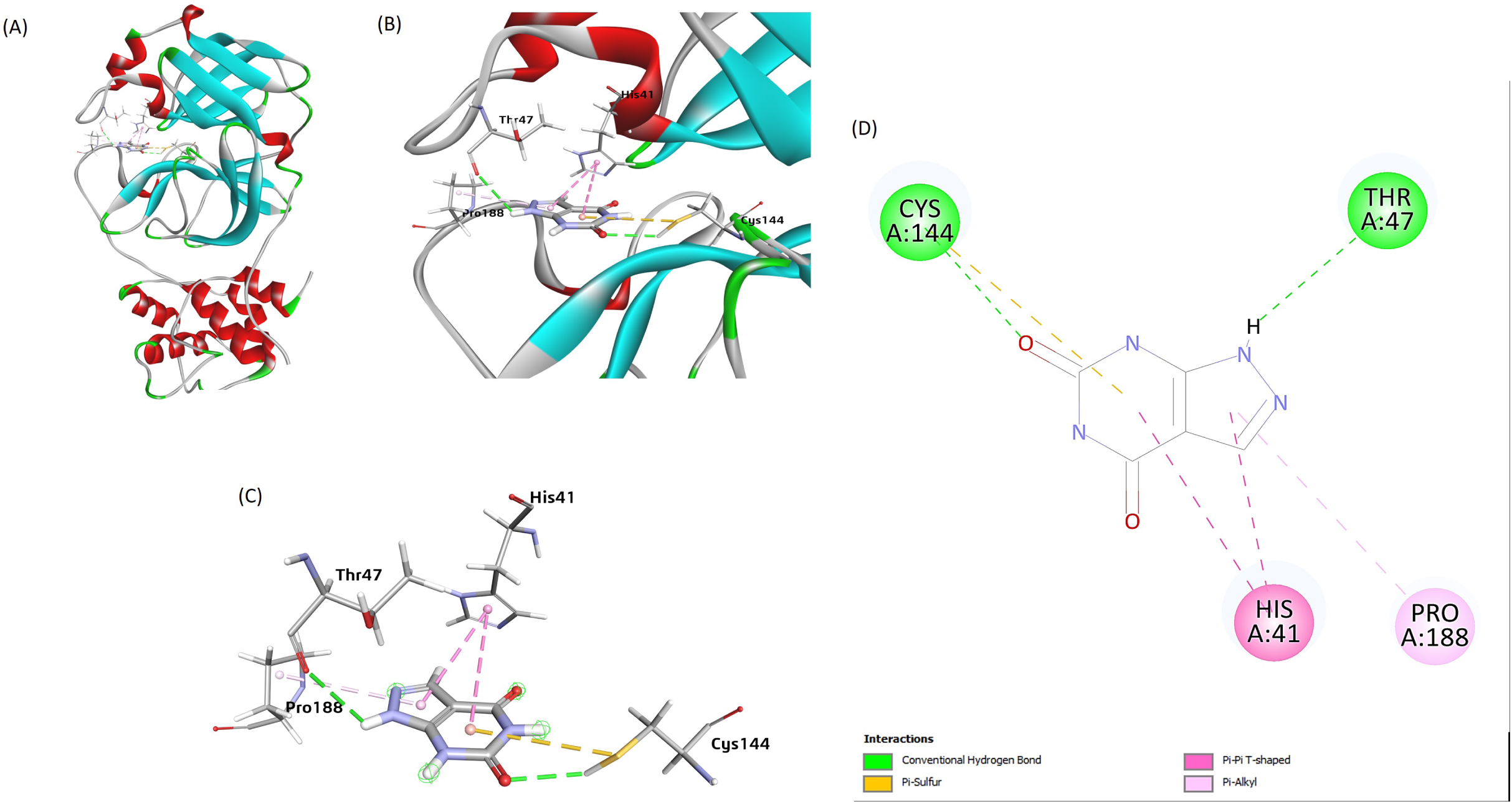
The best docking pose of the interaction of the FIPV-Mpro/ Oxipurinol interaction. (A) Mpro-Oxipurinol complex after docking. (B) and (C) showing Oxipurinol interaction with residues of the FIPV-Mpro. (D) The interaction of Oxipurinol with FIPV-Mpro residues, including (CYS144, HIS41, HIS162, THR47, and PRO188).

### 3.3. FIPV-Mpro interaction with the Favipiravir

The best docking pose for the interaction between the FIPV-Mpro and the **Favipiravir** shows a binding affinity with the highest binding energy (**ΔG**= **-115.58** Kcal/mol) (Table 1). The CDOCKER energy score for the same pose is (**-8.8)**. The interaction between the FIPV-Mpro and the **Favipiravir** involved some key amino acid residues and bonds, including (Glu-165 (Bond: hydrogen) and the Leu-164 (bond: Pi-alkyl), shown in the 2D image of (Figure 3).

**Figure 3:**
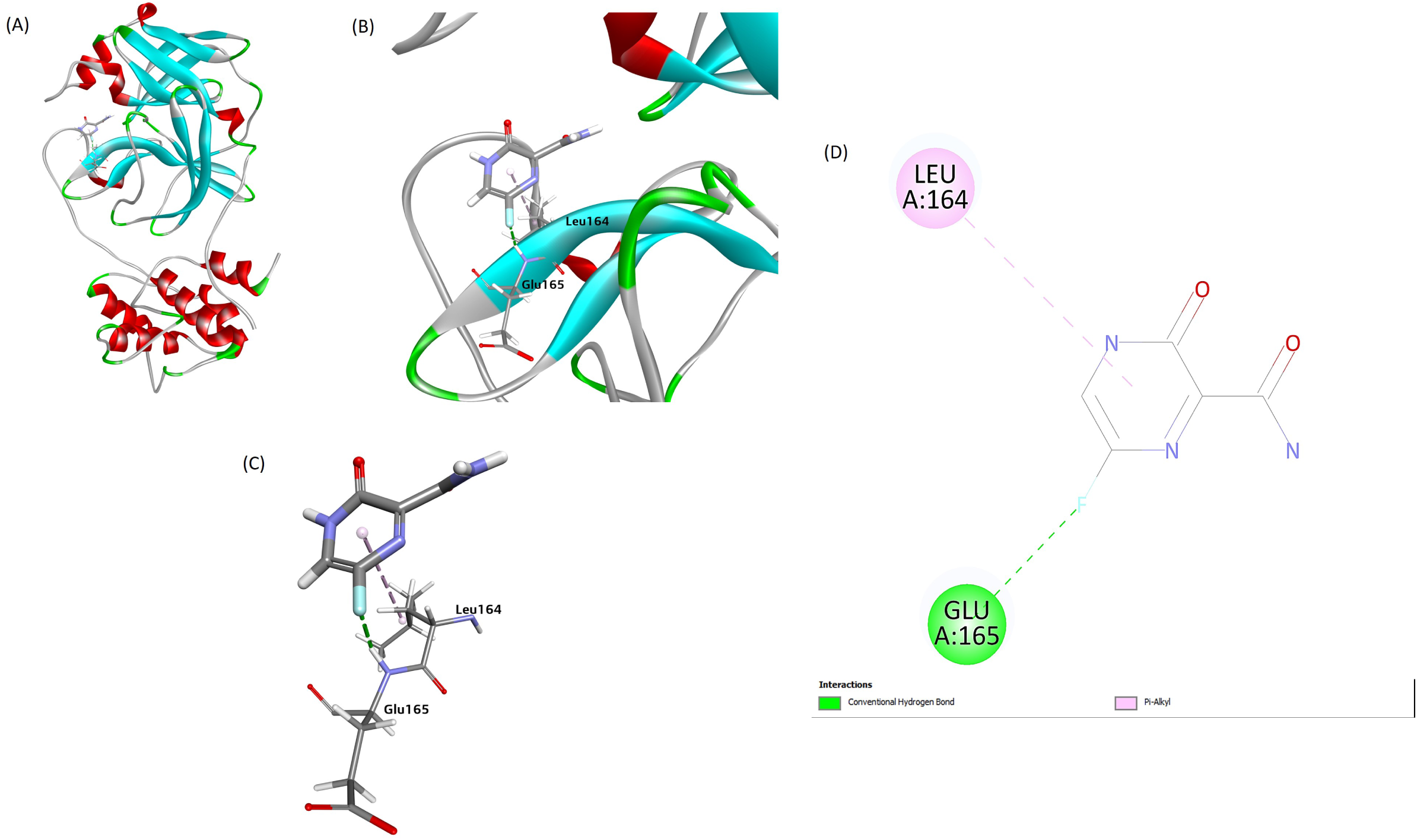
The best docking pose of the interaction of the FIPV-Mpro/Favipiravir interaction. (A) Mpro-favipiravir complex after docking. (B) and (C) showing favipiravir interaction with residues of Mpro. (D) Interaction of Favipiravir with FIPV-Mpro residues, including (LEU164 and GLU165).

### 3.4. FIPV-Mpro interaction with the pentoxifylline

The best docking poses docking for the FIPV-Mpro and the **pentoxifylline** showing a high binding affinity with the strongest binding energy (**ΔG**= **-130.82** Kcal/mol) (Table 1). The CDOCKER energy score for the same pose is (**-21.57)**. The interaction between FIPV-Mpro and the **Pentoxifylline** involved some key amino acid residues and bonds, including (Glu-165 (Bond: hydrogen) and the Leu-164 (bond: Alkyl), the His-41 (bond: Pi-Pi T shaped and Pi-Alkyl), the Pro-188 (bond: Pi-Alkyl)) as shown in 2D image (Figure 4) and (Table 1S).

**Figure 4:**
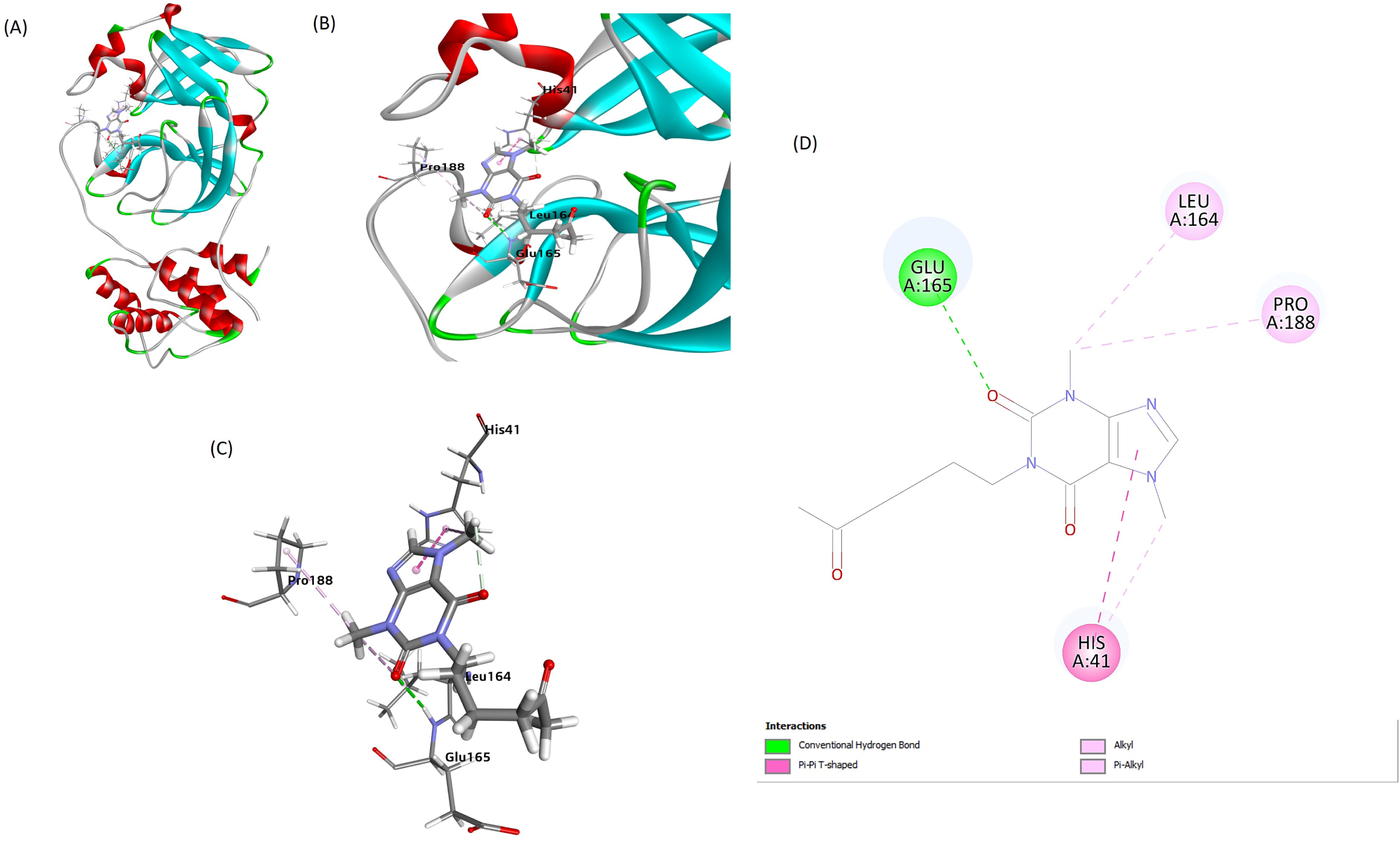
The best docking pose of the interaction of the FIPV-Mpro/Pentoxifylline interaction. (A) Mpro-pentoxifylline complex after docking. (B) and (C) showing pentoxifylline interaction with residues of Mpro. (D) Interaction of pentoxifylline with rFIPV-Mpro residues, including (HIS41, PRO188, LEU164, and GLU165).

### 3.5. FIPV-Mpro interaction with the Baricitinib

The best docking pose of FIPV-Mpro and the **Baricitinib** compound showed a high binding affinity with a strong binding energy (**ΔG**= **-162.05** Kcal/mol) (Table 1). The CDOCKER energy score for the same pose is (**25.79)**. The interaction between FIPV-Mpro and the **Baricitinib** compound involved some key amino acid residues and bonds, including (the Cys-144 (bond: Pi-Sulphur), the Thr-47 and the Phe-139 (bond: hydrogen), the Ala-141 (bond: Pi-alkyl), the His-41 (bond: hydrogen and Pi-sulfur), the Glu-165 (bond: Pi-Anion)) as shown in the 2D image (Figure 5) and (Table 1S).

**Figure 5:**
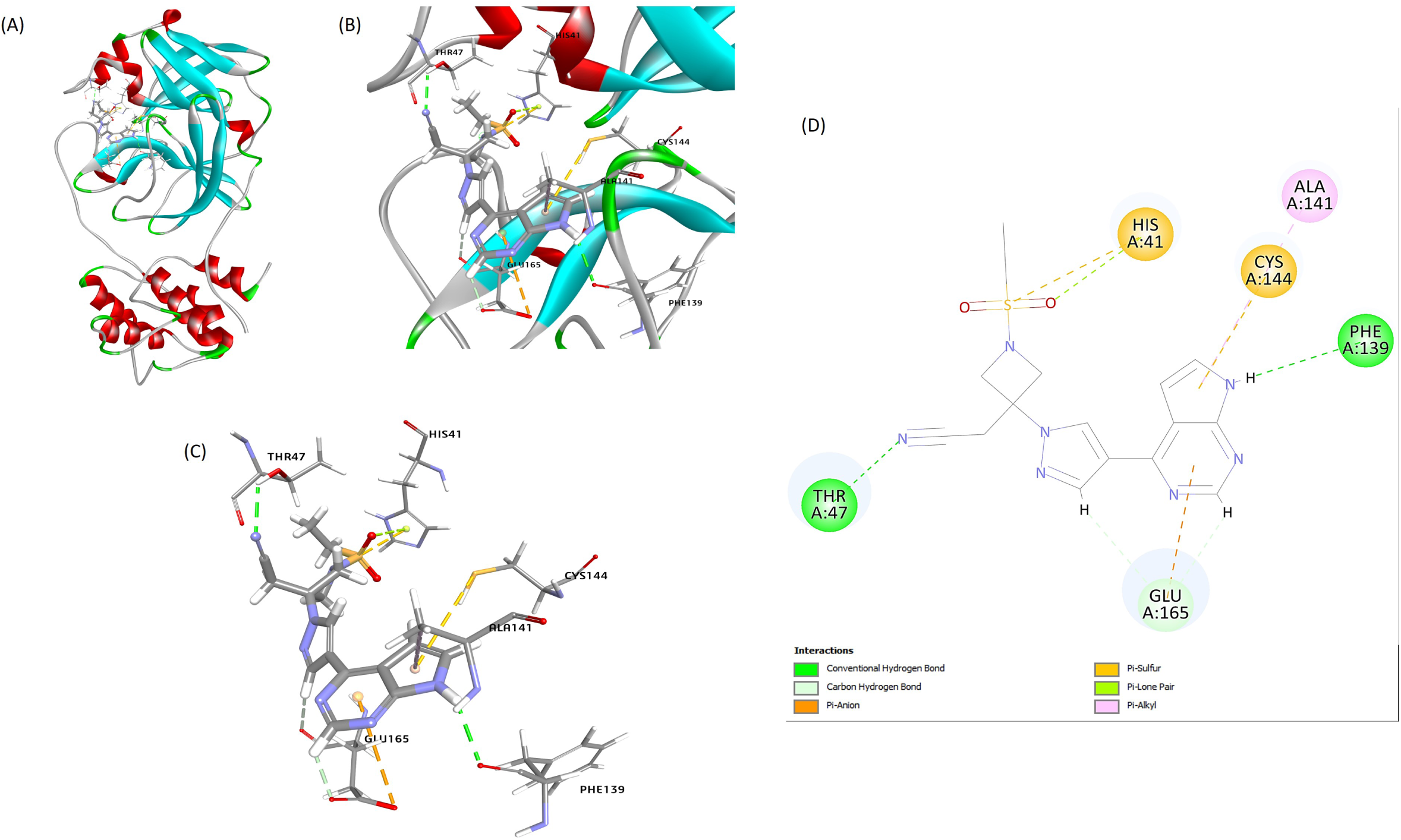
The best docking pose of the interaction of the FIPV-Mpro/Baricitinib interaction. (A) FIPV/Mpro-Baricitinib complex after docking. (B) and (C) showing Baricitinib interaction with residues of Mpro. (D) Interaction of Baricitinib with residues including (HIS41, ALA141, CYS144, PHE139, THR47, and GLU165).

### 3.6. FIPV-Mpro interaction with the methotrexate

The best docking pose of FIPV-Mpro and the **Methotrexate** shows the binding affinity with the highest binding energy (**ΔG**= **-229.42** Kcal/mol) (table 1). The CDOCKER energy score for the same pose is (**-44.55)**. The interaction between Mpro and **Methotrexate** involves key amino acid residues and bonds, including the Cys-144, the Leu-27and the Leu-164 (bond: Pi-Sulphur), the His-41, the Gln-187 and the Gly-167 (bond: conventional hydrogen and carbon-hydrogen), the Glu-165 and His-163 (bond: carbon-hydrogen), Val-26 (bond: unfavorable donor) shown in 2D image (Figure 6) and (Table 1S).

**Figure 6:**
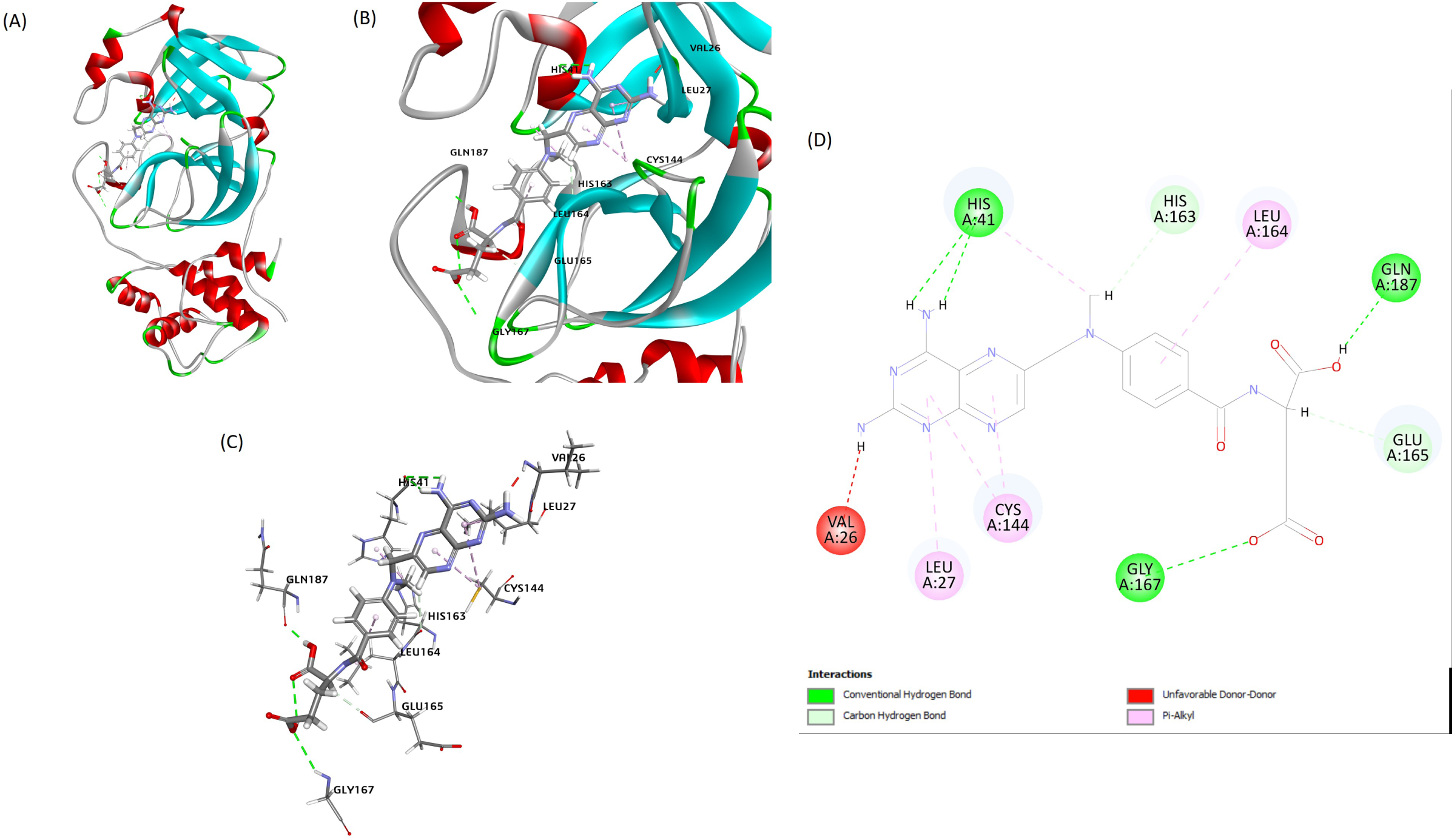
The best docking pose of the interaction of the FIPV-Mpro/Methotrexate interaction. (A) FIPV-Mpro-Methotrexate complex after docking. (B) and (C) showing Methotrexate interaction with residues of the FIPV-Mpro. (D) Interaction of Methotrexate with residues including (HIS41, HIS163, LEU164, GLN187, GLY167, CYS144, LEU27, VAL26 and GLU165).

### 3.7. FIPV-Mpro interaction with the Gemcitabine

The best docking pose of the **FIPV-Mpro-Gemcitabine** showed the binding affinity with the highest negative binding energy (**ΔG**= **-157.37** Kcal/mol) (Table 1). The CDOCKER energy score for the same pose was (**-9.6)**. The interaction between the **FIPV-Mpro-Gemcitabine** involved some key amino acid residues and bonds, including the Leu-164 (bond: carbon-hydrogen), the His-163, the Glu-165, and the Phe-139 (bond: hydrogen)) as shown in the 2D image (Figure 7) and (Table 1S).

**Figure 7:**
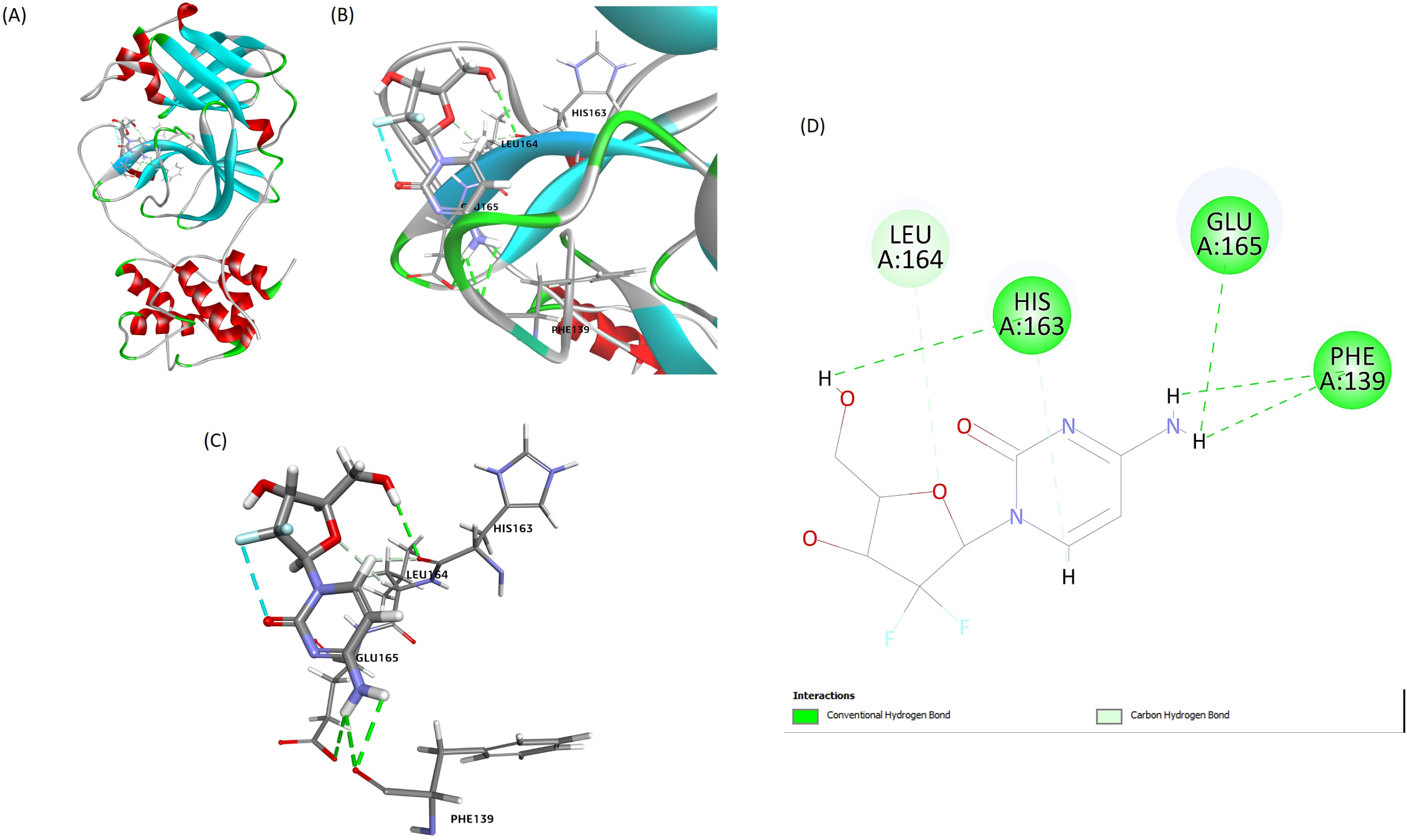
The best docking pose of the interaction of the FIPV-Mpro/Gemcitabine interaction. (A) the FIPV-Mpro-Gemcitabine complex after docking. (B) and (C) showing Gemcitabine interaction with residues of the FIPV-Mpro. (D) The interaction of the Gemcitabine with the FIPV-Mpro residues, including (GLU165, HIS163, LEU164, PHE139, and GLU165).

### 3.8. FIPV-Mpro interaction with the Galidesivir

The docking of the **FIPV-Mpro**-**Galidesivir** indicated the binding affinity with the highest negative binding energy (**ΔG**= **-201.36** Kcal/mol) (Table 1). The CDOCKER energy score for the same pose is (+**1.8)**. The interaction between the **FIPV-Mpro** and the **Galidesivir** involved some key amino acid residues and bonds, including the Pro-188 and the Thr-47 (bond: carbon-hydrogen), and the His-163 (bond: hydrogen)) as shown in the 2D image (Figure 8) and (Table 1S).

**Figure 8:**
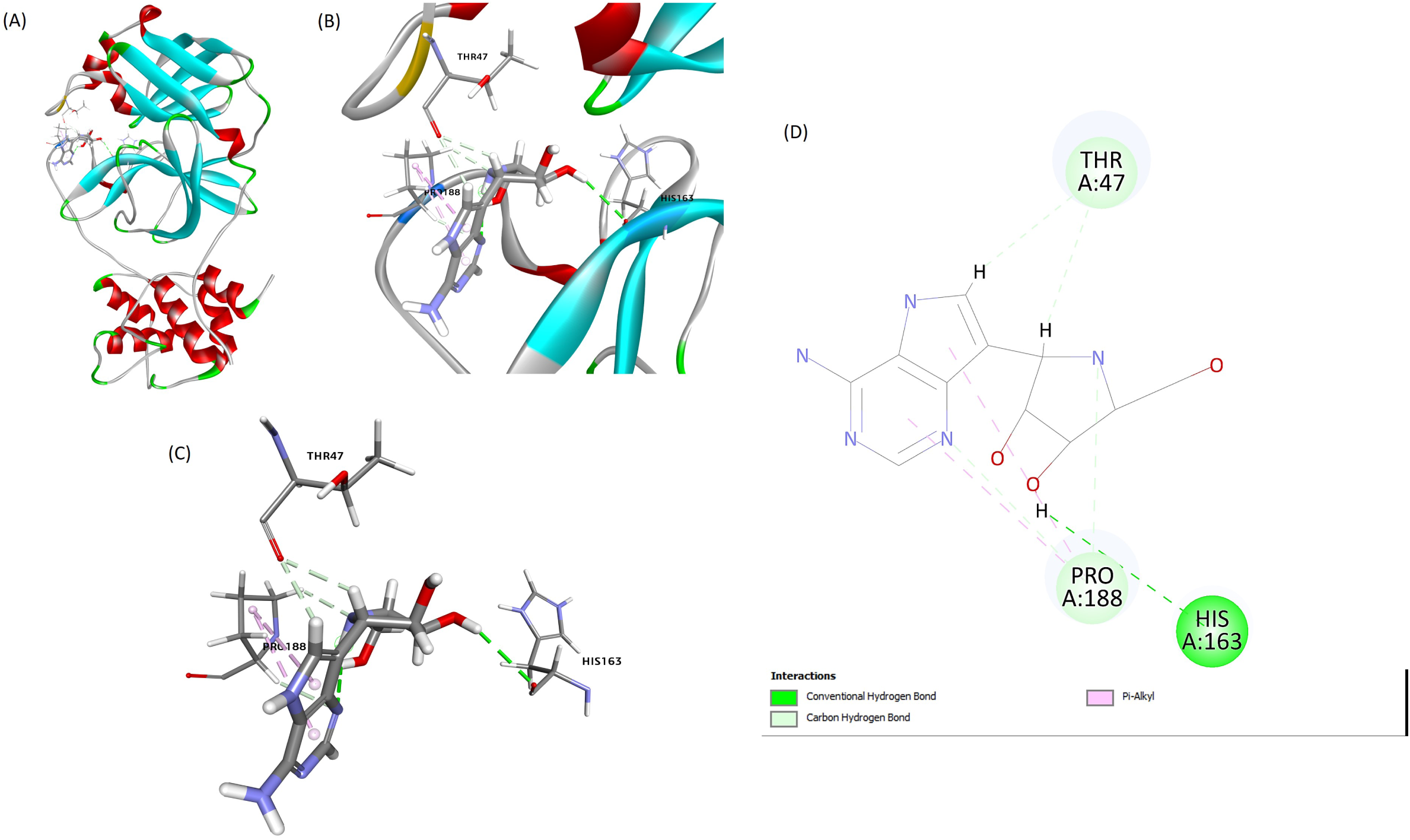
The best docking pose of the interaction of the FIPV-Mpro/Galidesivir interaction. (A) The FIPV/Mpro-Galidesivir complex after docking. (B) and (C) showing Galidesivir interaction with residues of the FIPV/Mpro. (D) Interaction of the Galidesivir with FIPV/Mpro residues, including (THR47, HIS163, and PRO188).

### 3.9. FIPV-Mpro interaction with the Ribavirin

The best docking pose of the **FIPV-Mpro interaction with the Ribavirin** indicated the binding affinity with the highest negative binding energy (**ΔG**= **-161.18** Kcal/mol) (Table 1). The CDOCKER energy score for the same pose was (+**15.23)**. The interaction between FIPV-Mpro and the **Ribavirin** involved some key amino acid residues and bonds, including the Pro-188 (bond: carbon-hydrogen), the His-163, Ser-189, and the Glu-165 (bond: hydrogen), the Leu-164 (Pi-Alkyl) showed in the 2D image (Figure 9 and Table 1S).

**Figure 9:**
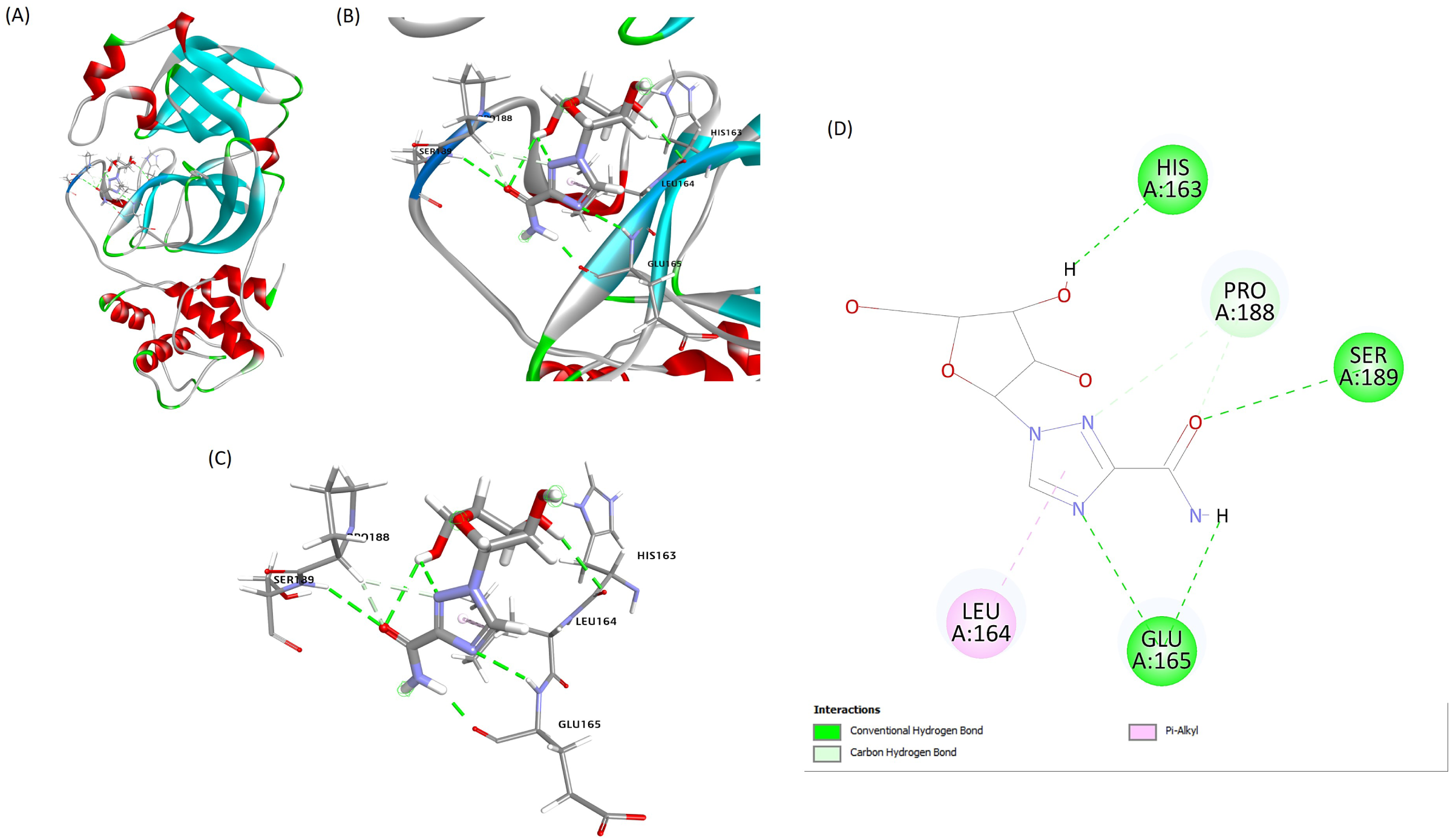
The best docking pose of the interaction of the FIPV-Mpro/Ribavirin interaction. (A) The FIPV-Mpro-Ribavirin complex after docking. (B) and (C) showing Ribavirin interaction with residues of the FIPV-Mpro. (D) The interaction of the Ribavirin with FIPV-Mpro residues, including (LEU164, GLU165, SER189, HIS163, and PRO188).

### 3.10. FIPV-Mpro interaction with the 6-Azauridine

The best docking pose of FIPV-Mpro **6-Azauridine** interaction confirmed the binding affinity with the highest binding energy (**ΔG**= **-148.33** Kcal/mol) (Table 1). The CDOCKER energy score for the same pose was (+**3.22)**. The interaction between Mpro and **6-Azauridine** involves key amino acid residues and bonds, including Pro-188 and His-163 (bond: carbon-hydrogen), the His-41 (bond: Pi-lone pair)) as shown in the 2D image (Figure 10) and (Table 1S).

**Figure 10:**
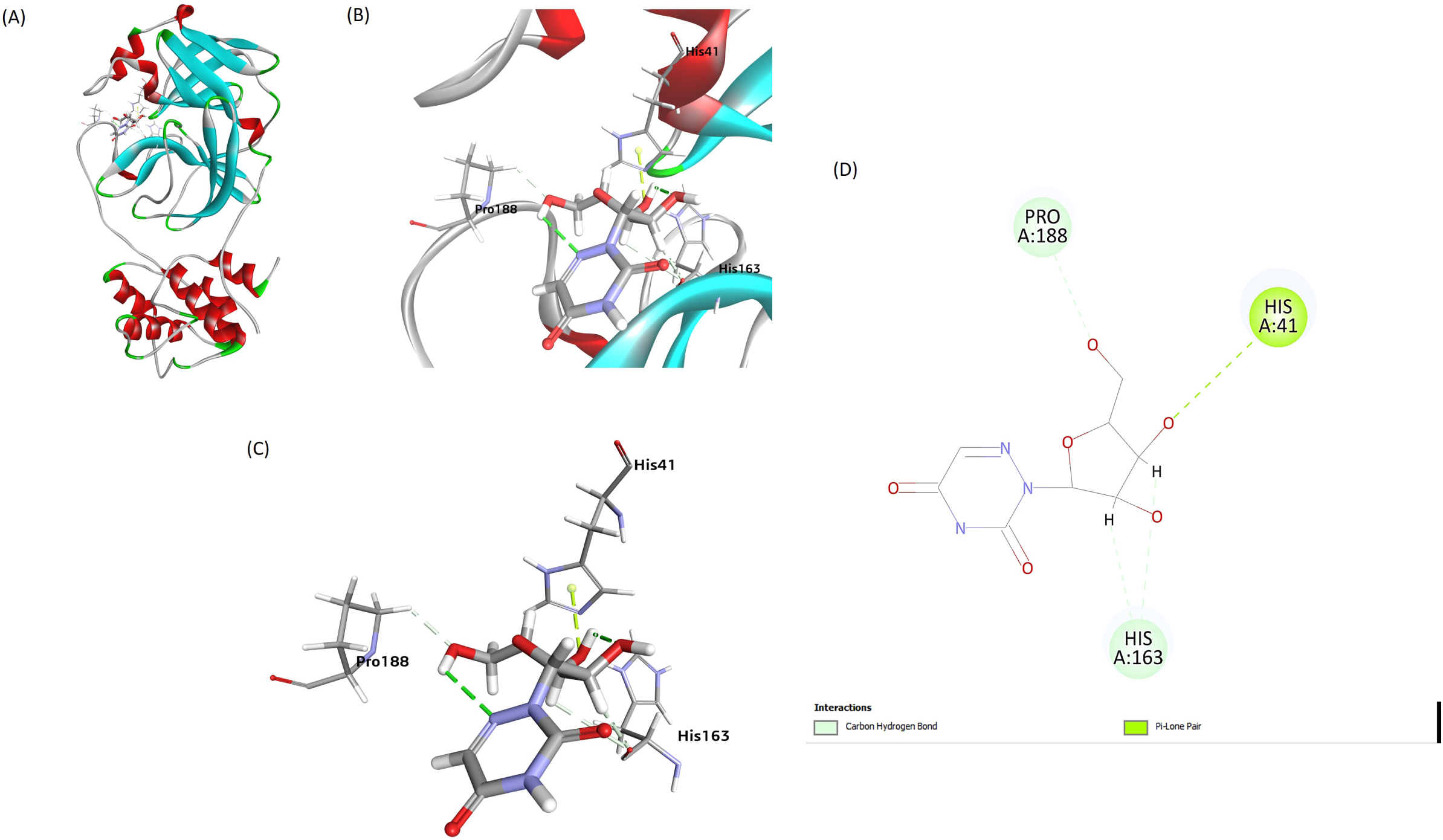
The best docking pose of the interaction of the FIPV-Mpro/6-Azauridine interaction. (A) The FIPV-Mpro-6-Azauridine complex after docking. (B) and (C) showing 6-Azauridine interaction with residues of Mpro. (D) The interaction of the 6-Azauridine with FIPV-Mpro residues, including (HIS41, HIS163, and PRO188).

### 3.11. FIPV-Mpro interaction with the GS441524

The best docking pose of the FIPV-Pro-**GS441524** interaction confirms the binding affinity with the highest binding energy (**ΔG**= **-151.54** Kcal/mol) (Table 1). The CDOCKER energy score for the same pose is (+ **4.08)**. The interaction between FIPV-Mpro and the **GS441524** involved some key amino acid residues and bonds, including the His-163 and the His-162 (bond: hydrogen), the His-41 (bond: Pi-Pi-T shaped), the Leu-164 (bond: Pi-alkyl)) as shown in the 2D image (Figure 11) and (Table 1S).

**Figure 11:**
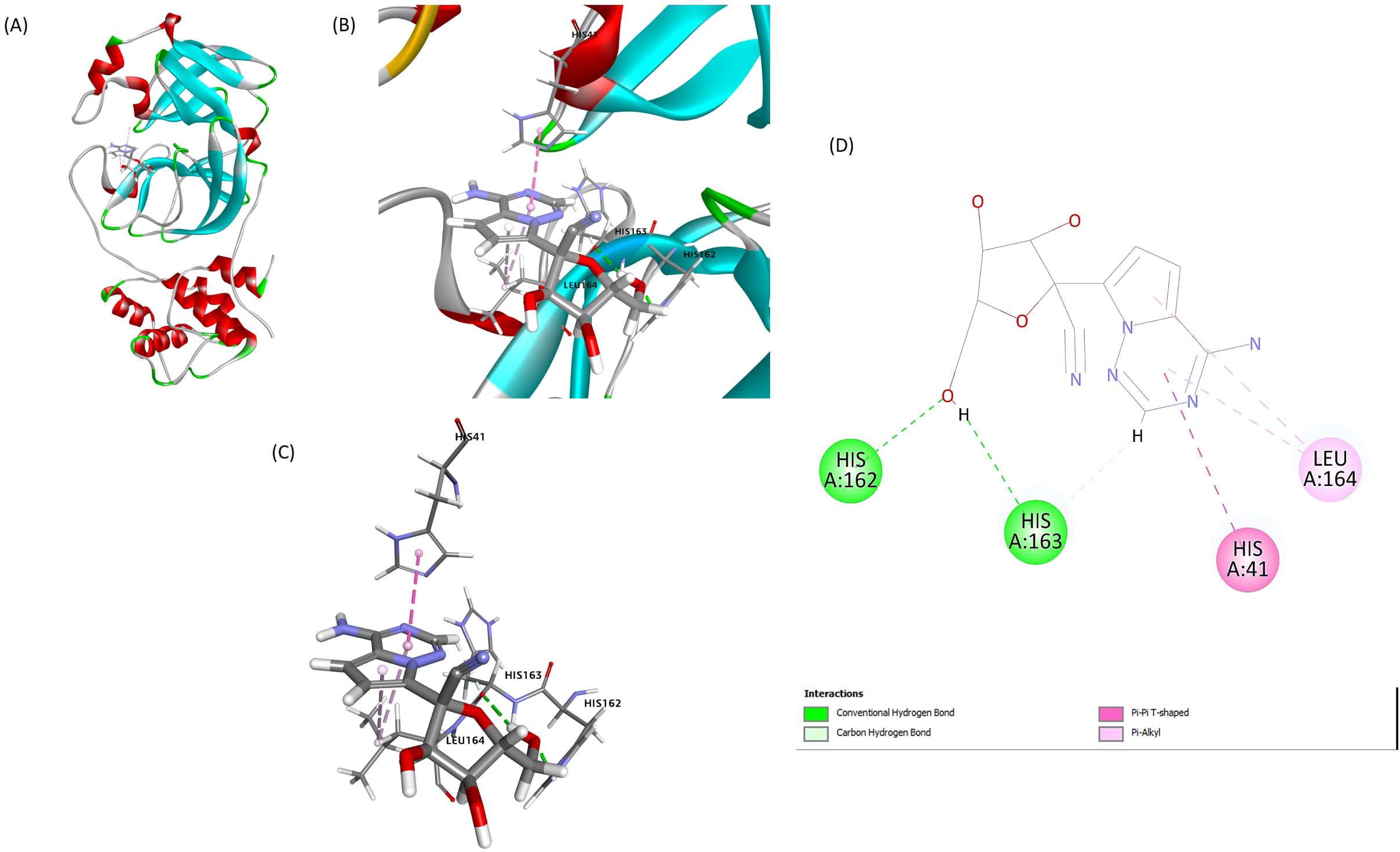
The best docking pose of the interaction of the FIPV-Mpro/GS441524 interaction. (A) The FIPV-Mpro-GS44152 complex after docking. (B) and (C) showing the GS44152 interaction with residues of FIPV-Mpro. (D) The interaction of the GS44152 with the FIPV-Mpro residues, including (HIS41, HIS162, HIS163, and LEU164).

### 3.12. FIPV-Mpro interaction with the Mizoribine

The best docking pose of the FIPV-Mpro-**Mizoribine** interaction confirmed the binding affinity with the highest negative binding energy (**ΔG**= **-152.54** Kcal/mol) (table 1). The CDOCKER energy score for the same pose was (**-7.10)**. The interaction between FIPV-Mpro and the **Mizoribine compound** involved some key amino acid residues and bonds, including the His-163 and the Ala-141 (bond: carbon-hydrogen), the Phe-139 (bond: hydrogen), the Cys-144 (bond: Pi-alkyl)) as shown in the 2D image (Figure 12) and (Table 1S).

**Figure 12:**
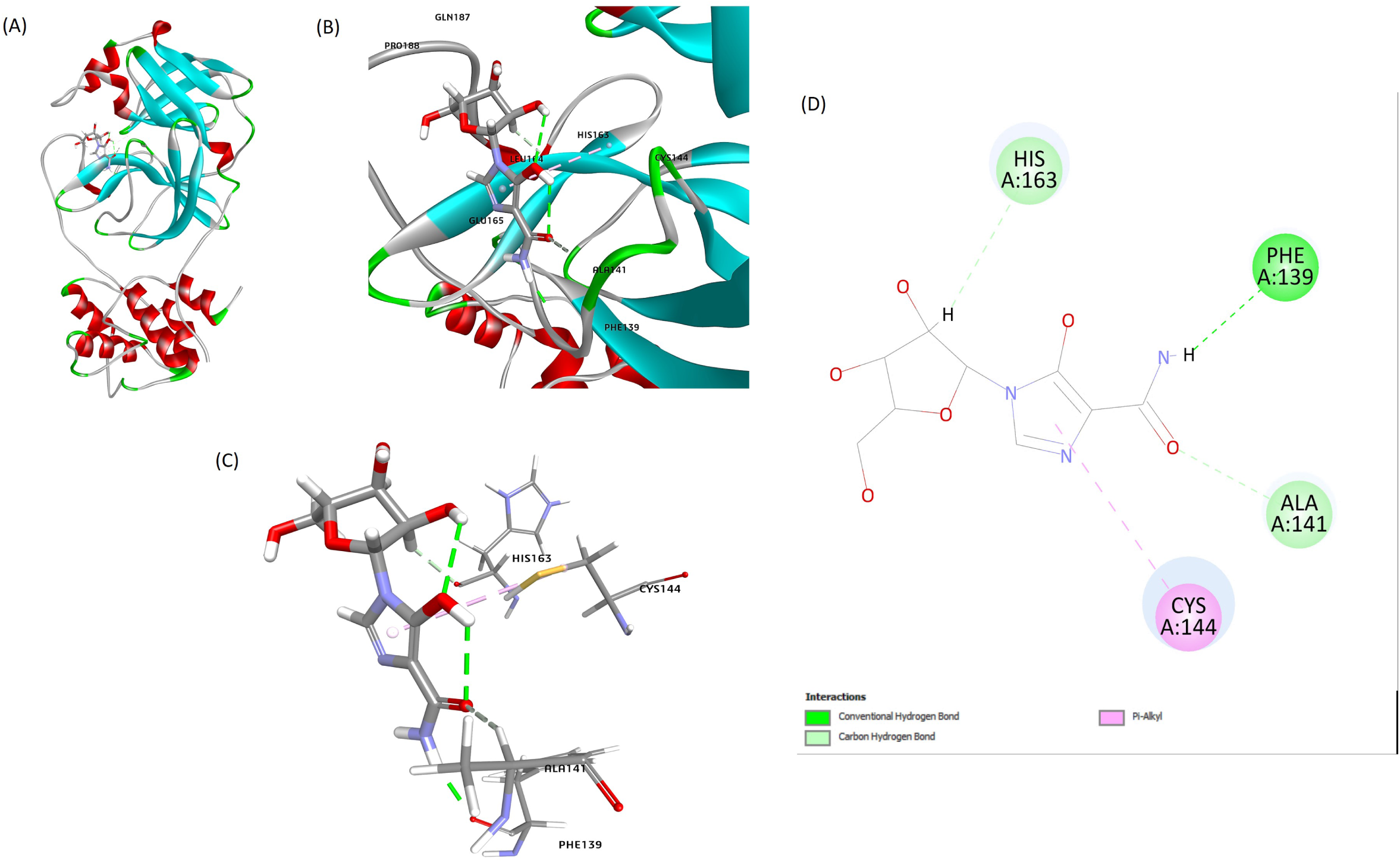
The best docking pose of the interaction of the FIPV-Mpro/Mizoribine interaction. (A) The FIPV-Mpro-Mizoribine complex after docking. (B) and (C) showing Mizoribine interaction with residues of the FIPV-Mpro. (D) The interaction of Mizoribine with the FIPV-Mpro residues, including (PHE139, ALA141, CYS144, and HIS163).

### 3.13. FIPV-Mpro interaction with the Sofosbuvir

The best docking pose of the FIPV-Mro-**Sofosbuvir** interaction confirmed the binding affinity with the highest binding energy (**ΔG**= **-232.76** Kcal/mol) (Table 1). The CDOCKER energy score for the same pose was (**-46)**. The interaction between the FIPV-Mpro and the **Sofosbuvir** compound involved some key amino acid residues and bonds, including the His-163 and the Thr-47 (bond: carbon-hydrogen), the Pro-188 (bond: Alkyl), the His-41 (bond: halogen)) as shown in the 2D image (Figure 13) and (Table 1S).

**Figure 13:**
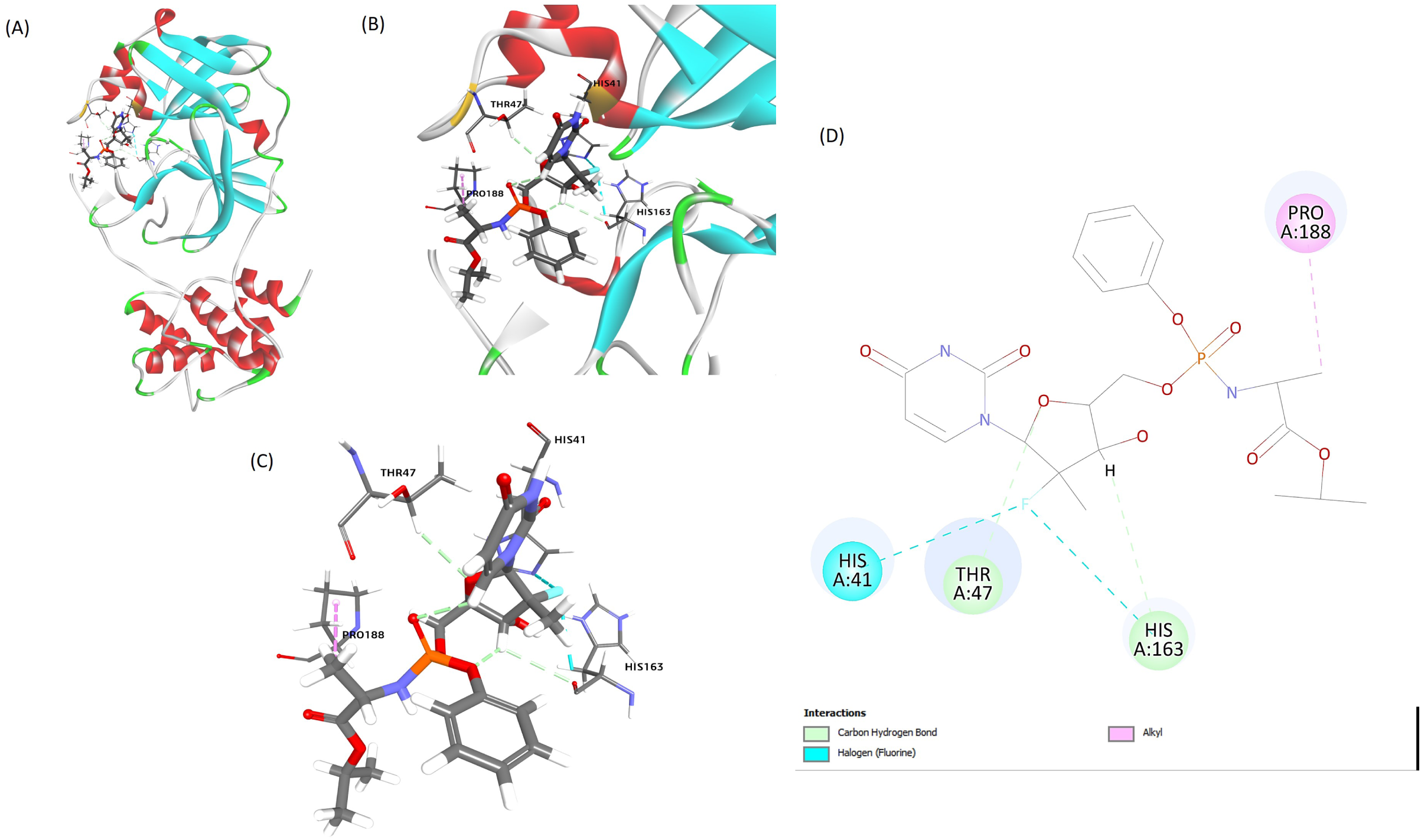
The best docking pose of the interaction of the FIPV-Mpro/Sofosbuvir interaction. (A) The FIPV-Mpro-Sofosbuvir complex after docking. (B) and (C) showing Sofosbuvir interaction with residues of the FIPV-Mpro. (D) The interaction of the Mizoribine with residues including (PHE139, ALA141, CYS144, and HIS163).

### 3.14. FIPV-Mpro interaction with the Molnupiravir

The best docking pose of the FIPV-Mpro-**Molnupiravir** interaction confirmed the binding affinity with the highest negative binding energy (**ΔG**= **-202.54** Kcal/mol) (Table 1). The CDOCKER energy score for the same pose is (**-12.96)**. The interaction between the FIPV-Mpro and the **Molnupiravir** compound involved some key amino acid residues and bonds, including the Leu-164 and the Thr-47 (bond: carbon-hydrogen), the Cys-144, the Leu-164 (bond: Alkyl and Pi-alkyl), the Glu-165 (bond: hydrogen), the Pro-188 (bond: Carbon-hydrogen)) as shown in the 2D image (Figure 14) and (Table 1S).

**Figure 14:**
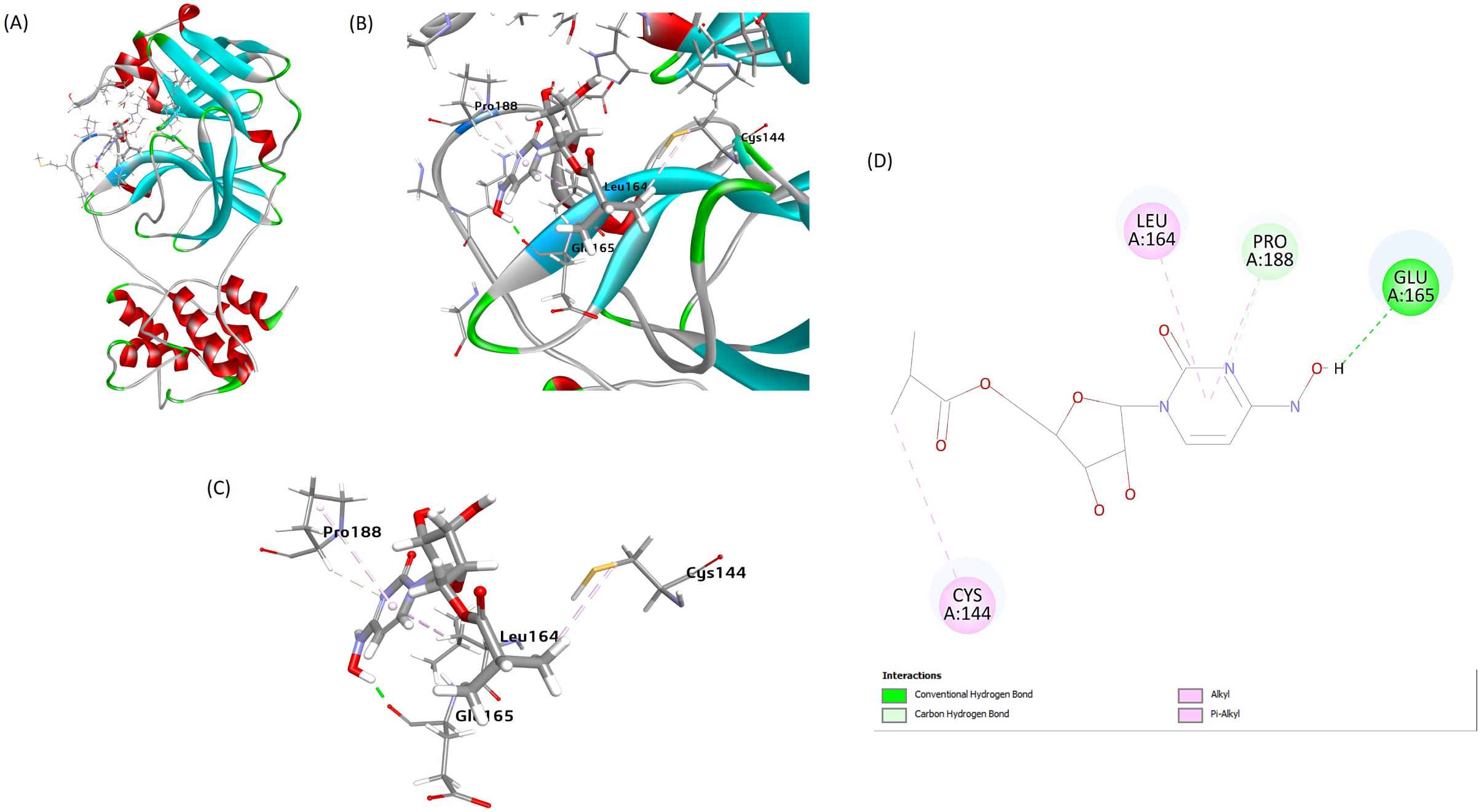
The best docking pose of the interaction of the FIPV-Mpro/Molnupiravir interaction. (A) The FIPV-Mpro-Molnupiravir complex after docking. (B) and (C) showing Sofosbuvir interaction with residues of the FIPV-Mpro. (D) The interaction of the Molnupiravir with the FIPV-Mpro residues, including (CYS144, LEU164, PRO188, and GLU165).

### 3.15. FIPV-Mpro interaction with the Tenofovir

The best docking pose of the FIPV-**Tenofovir** interaction confirmed the binding affinity with the highest binding energy (**-185.24** Kcal/mol) (Table 1). The CDOCKER energy score for the same pose was (**-30.53)**. The interaction between FIPV-Mpro and the **Tenofovir** compound involved some key amino acid residues and bonds, including the Leu-164 and the Thr-47 (bond: carbon-hydrogen), the Cys-144, the His-163, the His-162 and the Glu-165 (bond: hydrogen), the His-41 (bond: Pi-T shaped), the Cys-144 (bond: Pi-alkyl)) as shown in 2D image (Figure 15) and (Table 1S).

**Figure 15:**
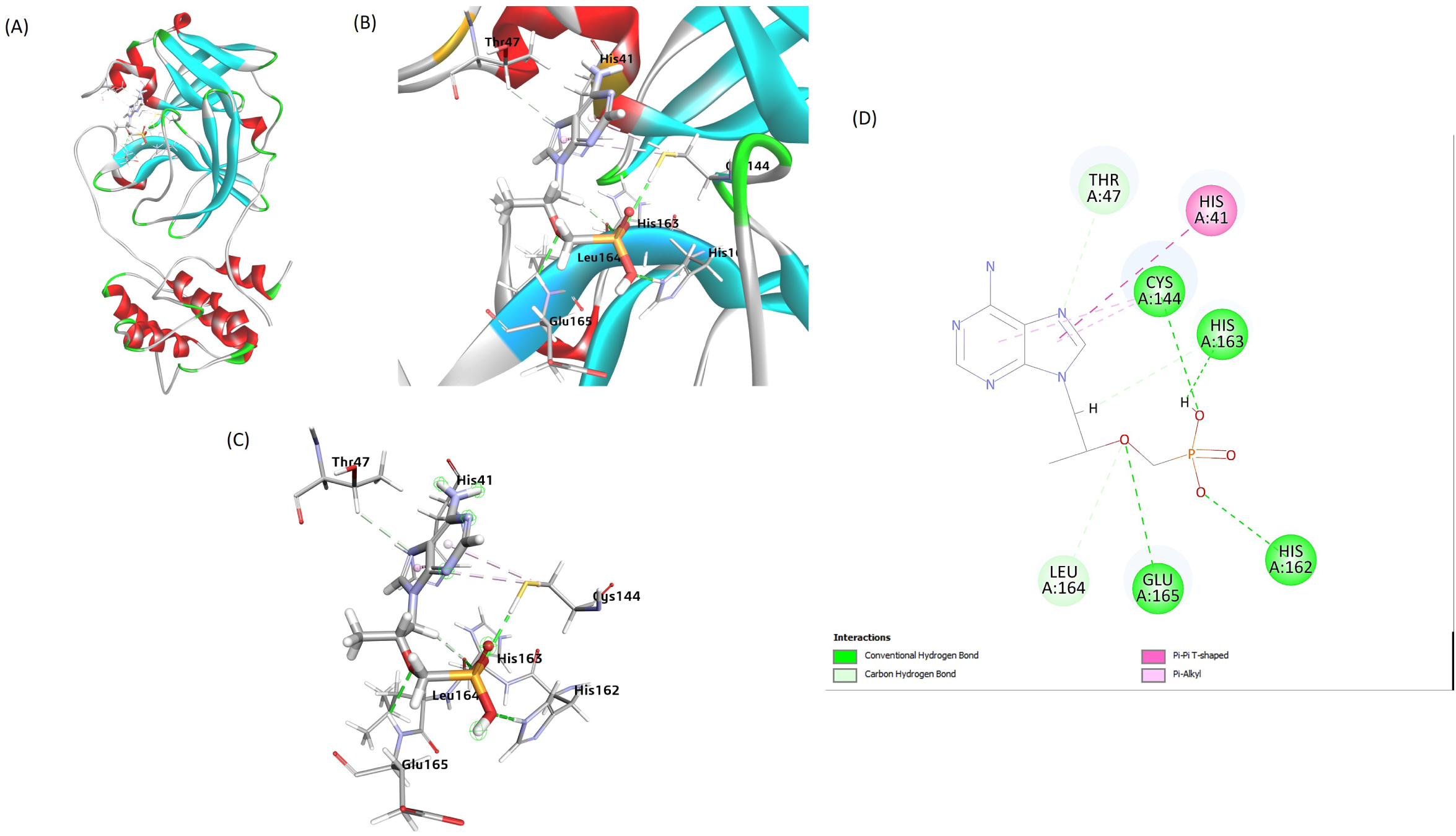
The best docking pose of the interaction of the FIPV-Mpro/Tenofovir interaction. (A) The FIPV-Mpro-Tenofovir complex after docking. (B) and (C) showing Tenofovir interaction with residues of Mpro. (D) The interaction of the Tenofovir with the FIPV-Mpro residues, including (HIS41, THR47 HIS162, HIS163, CYS144, LEU164, and GLU165).

## 4. Discussion

The FIPV hijacks the host cells’ transcriptional machinery and directs the infected cells to synthesize the viral proteins, particularly the two overlapping polyproteins, pp1a, and pp1ab, which cleave through coronavirus-encoded proteases—papain-like protease (PL^pro^) and the Mpro [5]. The cleavage of pp1a and pp1ab generates 16 non-structural proteins (Nsps), which play a role in viral replication [19]. Several compounds have been recently tried as antiviral therapy for FIPV infection in cats. Both the Remdesivir and the GC376 (protease inhibitor mainly targets the 3CLP) compounds showed impressive results as antiviral therapy against FIPV infection in cats using both the in vitro and in vivo models [6, 20]. The GS-441524 as nucleoside analogs showed a promising result as a potent antiviral compound against FIPV infection in vitro in cell culture as well as in vivo in cats [8]. Meanwhile, the oral administration of the GS-441524 showed more potent antiviral effects compared to the GC376 compound [21]. Among humans and animals, coronaviruses exhibit the presence of similar structurally related functional proteins, such as the Mpro, also known as 3CL protease (3CLpro), which plays a crucial role in viral replication [22]. The FIPV-Mpro features a catalytic dyad composed of two essential residues, His41 and Cys144, with the cysteine residue functioning as a nucleophile. This catalytic dyad is pivotal for the enzymatic activity of Mpro, facilitating the cleavage of viral polyproteins into functional proteins essential for viral replication and assembly [23, 24]. In the current study, the molecular docking of 15 selected compounds was conducted to identify some potential anti-FIPV agents from a pool of non-natural compounds. Using the CDOCKER for molecular docking, we evaluated and ranked the binding affinities of promising compounds with the catalytic dyad residues (His41 and Cys144) in the binding pocket of FIPV-Mpro. The binding affinity CDOCKER scores of the top three ranked compounds and Mpro structure ranged from (−46.7 to −12.2), (Table 1). These three top-ranked compounds were selected based on the calculated binding energy (ΔG) using Biovia Discovery Studio. The binding energy scores of the top three-ranked complexes between ligands and the Mpro ranged (-232.76 to -202.54) kcal/mol (Table 1). Out of 15 selected compounds, the top three compounds that exhibited the best binding affinities in the catalytic dyad pockets are (Sofosbuvir, Methotrexate, and Molnupiravir) (Table 1). The CDOCKER and binding energy scores of reference ligand N3 with Mpro in the same binding pocket were (-78 and -246 kcal/mol). The Michael acceptor inhibitor (N3) compound showed a marked inhibitory effect on the SARS-CoV-2 Mpro compared to other compounds [24]. Our in silico and molecular docking results of the N3 compound on the FIMPV-Prop are very consistent with that of the SARS-CoV-2 (Figure 1 and Tabel 1). The binding energy of all the compounds, including the reference ligand (N3) with FIPV-Mpro, is shown in (Figure 16). **Sofosbuvir is a nucleoside analog currently used clinically and is an** approved drug used to treat HCV infection in humans. Sofosbuvir is a prodrug that is hydrolyzed by liver enzymes after absorption to form the monophosphate uridine analog, which is further phosphorylated to form the active triphosphate form [25]. **The Sofosbuvir** has been recently repurposed to inhibit the SARS-CoV-2 replication by targeting the viral RNA-dependent RNA polymerase, disrupting the viral life cycle [26, 27]. Sofosbuvir-based treatment regimens could potentially reduce the case fatality rates of severe SARS-CoV-2 infection and reduce the associated complications of the viral infection in humans [28]. In our study, Sofosbuvir interacts with the FIPV-Mpro-HIS41 residue (halogen bond) of the catalytic dyad (Figure 13 and Table 1S), and this interaction may potentially inhibit its binding with FIPV-PP1ab. Methotrexate (MTX) is a Dihydrofolate reductase inhibitor that acts mainly by suppressing severe inflammatory reactions, particularly following SARS-CoV-2 infection. In a similar pattern, FIPV induces severe inflammatory reactions through the induction of immune-mediated coronaviral vasculitis [29]. A recent study showed that methotrexate targets SARS-CoV-2 specific proteins and enzymes by inhibiting diverse protein families such as proteases, reductases, synthases, and transformylase [30]. The inhibitory action of methotrexate of these enzymes leads to inhibition of virus entry and virus replication [31]. In our study, methotrexate interacts with the FIPV-Mpro-HIS41 residue (halogen bond) and CYS144 (Pi-Sulphur bond) of the catalytic dyad (Figure 6) and (Table 1S), and this interaction may inhibit its binding with FIPV-polyproteins, pp1a and pp1ab. Molnupiravir is a nucleoside analog used as an antiviral drug against the influenza virus and SARS-CoV-2 infection [32]. Although all three ligands occupied the same binding site, the interaction of Molnupiravir was found to be better than that of sofosbuvir due to the higher hydrogen bond interactions with the FIPV-Mpro. In terms of binding energy, methotrexate and sofosbuvir show better binding, and Molnupiravir shows lower binding energy than sofosbuvir. In the current study, the Molnupiravir interacts with the FIPV-Mpro CYS144 (Alkyl and Pi-alkyl bond) (Figure 14) and (Table 1S) of the catalytic dyad, and this interaction may inhibit its binding with polyproteins, pp1a, and pp1ab. The FIPV-Mpro-mediated cleavage of these proteins is responsible for virus replication inside the host cell. Our docking study suggests repurposing some antiviral agent targeting the FIPV-Mpro, either as a sole drug or in combination with other immune-modulatory compounds.

**Figure 16:** The histogram showing the binding energy of the selected compounds’ interaction with the FIPV-Mpro in a descending order compared to the N3 as a reference ligand

## Data availability statement

All the data is available upon request from the corresponding author.

